# Selective Vulnerability of Senescent Glioblastoma Cells to Bcl-XL Inhibition

**DOI:** 10.1101/2020.06.03.132712

**Authors:** Masum Rahman, Ian Olson, Moustafa Mansour, Lucas P. Carlstrom, Rujapope Sutiwisesak, Rehan Saber, Karishma Rajani, Arthur E Warrington, Adam Howard, Mark Schroeder, Sisi Chen, Paul A. Decker, Eliot F. Sananikone, Yi Zhu, Ian F. Parney, Sandeep Burma, Desmond Brown, Moses Rodriguez, Jann N. Sarkaria, James Kirkland, Terry C. Burns

## Abstract

Despite decades of research and numerous basic science advances, there have only marginal gains in improving glioblastoma multiforme survival. Therefore, new ideas and approaches for treating this aggressive disease are essential to drive progress forward. Conventional therapies, such as radiation and Temozolomide (TMZ), function to cause oxidative stress and DNA damage yielding a senescent-like state of replicative arrest in susceptible cells. However, increasing evidence demonstrates malignant cells can escape senescence leading to tumor recurrence. Ablation of non-replicating senescent tumor cells after radiation and chemotherapy may be an avenue to reduce the rates of tumor recurrence. Senolytic agents have been developed that selectively target senescent cells, but it remains unknown whether senolytics might be utilized against senescent-like glioma cells. We employed radiation or TMZ to induce a functionally senescent state in human glioblastoma cells. Viable cells that survive these treatments were then utilized to screen candidate senolytic drugs, to identify those selectively effective at ablating senescent-like cells over naïve non-tumor and proliferative cells. Among 10 candidate senolytic drugs evaluated, only Bcl-XL inhibitors demonstrated reproducible senolytic activity in radiated or TMZ-treated glioma across the majority of GBM cell lines evaluated. Conversely, Bcl-2 inhibitors and other established senolytic drugs failed to show any consistent senolytic activity. In agreement with these data, Bcl-XL knockdown selectively reduced the viability of senescent-like GBM cells, whereas knockdown of Bcl-2 or Bcl-W yielded no senolytic effect. These findings demonstrate the potential to harness radiation-induced biology to ablate latent surviving cells and highlight Bcl-XL dependency as a potential vulnerability of surviving GBM cells after exposure to radiation or TMZ.

## Introduction

Gliomas are fatal infiltrative brain tumors for which standard treatment includes maximal surgical resection followed by radiation and Temozolomide (TMZ).[1] Treatment-resistant cells may persist in a latent state for months or years prior to inevitable recurrence. Since surgery is rarely performed until radiographic or symptomatic recurrence, relatively little is known about the molecular characteristics and potential drug sensitivity of the latent human glioma cells that ultimately give rise to recurrence.[2,3,4] Prior work has demonstrated that radiation and TMZ induce a reversible senescent-like phenotype in malignant glioma.[2][5] Senescent cells are characterized by altered transcriptional, metabolic and secretory phenotype all while disconnected from the cell cycle. Although radiation and cytotoxic chemotherapy can induce a senescent-like state of cell cycle arrest in cancer, malignant cells have been shown to escape senescence and re-enter cell cycle, leading to refractory tumor recurrence. [6][7] Recent advances in senescent cell biology have been fueled by evidence that senescent cells underlie mechanisms of aging and aging-associated disease. As such, a growing array of drugs has been identified to ablate senescent cells. Senolytic drugs identified to date target several mechanisms implicated in cancer. Fortunately, many such “senolytic” drugs were originally developed as therapeutics for oncology with toxicity profiles already established in human clinical trials and provide opportunities for more rapid application in other fields where their effects may provide benefit. Since conventional therapy with TMZ and radiation frequently induces a latent period of non-proliferation prior to inevitable recurrence, we investigated whether known senolytic drugs target human GBM cells surviving in a non-proliferative senescent-like state. Herein, we demonstrate that post-chemotherapy or radiation treated GBM cells are selectively vulnerable to BCL-XL inhibition.

## Materials and methods

### Cell culture

Human patient-derived glioblastoma lines utilized have been previously described and were cultured according to established protocols.[8] Tumor lines maintained as patient-derived xenografts (PDX) are from the National PDX resource.[8] Such lines are designated as “GBM-6,10,12,39,76,123,164,196”. Protocols for implantation of patient-derived Glioblastoma cells, serial passage of flank tumor xenografts, and short-term explant culturing have described previously,[8] with some lines maintained in serum-containing media, and some lines maintained in serum free media as noted in table-2). Some cell lines were maintained from time of harvest as in vitro cell lines rather than PDX lines, as previously reported protocol.[9] Such lines are designated as dBT114,116,120,132(differentiated brain tumor). Culture conditions were unchanged after induction of senescence.

### Senescence induction

After plating, cells were maintained in 10cm culture dishes for 2-4 days until >50% confluent. They were then treated with TMZ for 7days or radiation (cesium gamma radiator) with varying doses as indicated.[5][10] Most experiments were performed after use of 15Gy to induce senescence. Radiation and TMZ each caused death of a variable percentage of cells in the ensuing days. “Senescent” tumor cells used for experiments are designated by those that survive following TMZ or radiation treatments. Except for radiation doses of 4Gy or below, no visible proliferation occurred within 1 month after treatment with TMZ or radiation. For initial screening of senolytic drugs, cells were maintained for at least 20 days after radiation or TMZ prior to re-plating cells into black walled optical 96-well plates (5000cells/well) for doing drug treatment.

### Analysis of cell proliferation

0Gy,4Gy, 8Gy, 15Gy, and 20Gy radiation was delivered as described above. Three days thereafter, cells then plated in 96-well plates. After allowing cells to adhere overnight, they were then placed into the IncuCyte (Thermo Scientific Series 8000 WJ Incubator) and images were captured every four hours for automated quantification of cellular confluence.

### Senolytic drug screen

Human glioblastoma cells were exposed to 10,15 or 20Gy radiation as indicated above. All GBM lines tested (Table-2) yielded a subpopulation of surviving non-proliferative cells used for subsequent experiments, with the exception of GM43 which yielded insufficient surviving cells for senescence experiments. All senolytic drugs used in this study were dissolved in DMSO (Table-1). Control (0nM) cells were treated with the same dose of DMSO as cells with the highest drug concentration. Unless specified otherwise, cells were maintained in drug-containing media for four days prior to evaluation of cell viability using ATP lite, Cell-Titer-Glo assay according to company protocol (Cat# G7570). For all experiments, luminescence values are normalized to the 0nM control for that cell line and radiation dose. All dose response graphs are depicted standard error of the mean (SEM) of 3 technical replicates at each concentration. Representative data are shown for experiments performed in independent replicates; complete normalized data for all assays performed (>10,000) are provided in online supplemental data.

### Evaluation of radiation dose, drug exposure time dependency

To evaluate the radiation dose effect on BCL-XL inhibitor sensitivity, we radiated GBM39 with 0Gy,1Gy,2Gy,4Gy,8Gy,15Gy. Four days after radiation, cells were plated in black wall 96wells plate. Overnight and drugs added the next day. Seven days after drug treatment, cell viability was measured by Cell Titer-Glo. To determine the impact of varying duration of drug exposure, we plated 15Gy radiated cells 7days after radiation, where the minimum drug exposure time was 1hour and the maximum 96hours. Cells were plated in black-wall 96-well plates, and drugs added the next day as described above. At the designated time-point, drug-containing media was removed and replaced with drug-free media for the remainder of the experiment (one wash with media, during a replacement). The cell viability assay was performed at the end of 96 hours.

### qRT-PCR

RNA was extracted from cells previously radiated at 15Gy at 1, 3 and 7 days. Briefly, cells were washed with PBS before being homogenized with TriZol reagent (Invitrogen). RNA precipitation was performed at −20°C overnight. Resulting RNA pellets were dissolved in RNase-free water and concentration was measured by absorbance at 260 nm (A260) using Nanodrop2000. cDNA synthesis was performed with 1 ug of total RNA using M-MLV reverse transcriptase kit (ThermoFisher) as per the manufacturer’s protocol. 25 ng of cDNA was used for real-time PCR with Taqman gene expression assay targeting IL-6 (Hs00174131_m1), Bcl-2 (Hs00608023_m1), Bcl-XL (Hs001691412_m1) on ABI 7500/7500-Fast Real-Time PCR System (Applied Bioscience). The relative expression of each gene was determined by the ΔΔCT method.

### SA-β-Gal staining

Senescence-associated β-galactosidase staining Kit (Cell Signaling Technology #9860) was used as an indicator of relative senescence after radiation as per the manufacturer’s directions. Briefly, cells were fixed for 10 min in β-galactosidase fixative Solution (10% 100x Fixative Solution; 90% H2O), and washed with PBS. The cells were then stained with β-Galactosidase Staining Solution (93% 1x Staining Solution; 1% 100x Solution A, 1% 100x Solution B, 5% X-gal. Wells without samples were filled with PBS, and the plate was wrapped in parafilm to prevent evaporation. The plate was left overnight in a dry incubator at 37° C. The next day, cells were examined under a microscope for β-gal-positive cells (blue staining). PBS was added to the wells, and the plate was placed on the rocker (speed = 30/min) for 5 minutes. The plate was washed three times. Staining was performed in sham and vehicle-treated control, 10Gy, 15Gy radiated, and TMZ treated GBM39 cells. To see the senescence maturation over time, Beta-Gal staining performed at day 0, 7, and 14 following 15Gy radiation. For each time points and conditions staining done at multiple wells.

### Protein analysis by western blotting

Cells grown in six-well plates, 10 cm dishes or T-25 flasks were washed with PBS, trypsinized and collected in 1.5 micro-centrifuge tubes as a cell pellet. The cell pellet was then lysed using lysis buffer composed of 10% RIPA lysis buffer, 4% Protease-Inhibitor cocktail, 1% Phosphatase-Inhibitor cocktail-2, 1% Phosphatase-Inhibitor cocktail-3, and 84% Molecular grade water. The cell palate with the lysis buffer then sonicated for 30 minutes in a water bath sonicator (one minute sonication every other minute for a total of 30 minutes). The whole lysate was centrifuged for 10 minutes at the speed of 17,000g to collect the supernatant as the final protein lysate. The concentration of the final protein lysate was then measured using the BCA kit (ThermoFischer). Proteins extracted from cells using lysis buffer, were separated in an SDS-polyacrylamide gel electrophoresis along with the protein ladder (Life Technology) using 4-12% Bis-Tris gel (ThermoFischer). Proteins were then transferred to a polyvinylidene difluoride (PVDF) membrane (Bio-Rad). The membrane was blocked in 5% fat-free milk (Cell-signaling technology) for 30 minutes, washed three times (5 minutes each) using Tris-buffered saline with tween20 (TBST) and probed with different antibodies (Cell-signaling technology).

### Gene knock-down using siRNA

siRNA sequences were either designed manually using commercially available soft wares or purchased as an already designed sequences (Horizon-ThermoFisher). The siRNA was re-suspended in 1X siRNA buffer (Dharmacon) or in any other Nuclease-free solutions. Cells were plated overnight with the optimal density in six-well plates (4×10^5^ per well) or 96-well plates (2×10^4^ per well) in antibiotic-free medium. The next day, the transfection complex was prepared by mixing the siRNA (either for the gene of interest or the negative siRNA control) and Lipofectamine RNAiMAX (Invitrogen) in serum-free medium, 250uL from the transfection complex were added to each well of the six-well plate and 10μL for each well in 96-well plate. One day post transfection RNA was collected to be analyzed by qRT-PCR to confirm gene silencing. 2-3 days post transfection, protein was collected to be analyzed by the SDS-page and western blotting to confirm gene silencing. Three days post transfection, the cell viability in response gene silencing was measured using Cell Titer-Glo assay (Promega), and the cell survival ratio was calculated compared to the negatively silenced control cells.

### Statistical analysis

Multiple linear regressions were used to assess the relationship of drug dose and treatment (e.g. control, radiation, TMZ) with cell inhibition within each cell line. For these models, cell inhibition was the outcome and drug dose, treatment, and the drug dose by treatment interaction were the independent predictors. If the drug dose by treatment interaction was significant (p<0.05), pairwise comparisons were made at each drug dose between treatment groups using a two-sample t-test. Statistical analysis was performed in Microsoft Excel. IC50 was calculated by nonlinear regression (Curve fit) of dose-response inhibition curve by using GraphPad prism 8.4.2.

## Results

Chemoradiation induces a senescence-like state of sustained proliferative arrest. We sought to test the hypothesis that glioblastoma could be relatively more sensitive to senolytic ablative therapies when induced into a senescent-like state of proliferative arrest following radiation.[10][11] No phenotypic marker is perfectly sensitive or specific for senescence. Indeed, significant overlap exists between phenotypic characteristics of malignant and senescent cells.[12][13] However, while malignant cells are defined in part by proliferative behavior, senescent cells, by definition, do not divide. We evaluated single fraction radiation doses in human glioblastoma cell lines and observed sustained loss of culture expansion with 8Gy or higher radiation in the GBM39 cell line (Figure 1). Senescent-like cultures demonstrated evidence of increased SA-β gal staining; consistent with a senescent phenotype induced by radiation (Figure 1A, Supp Fig-7A-C).[10] Higher expression of the senescence-associated gene, p21, and the anti-apoptotic pathway up-regulation, was also apparent in residual cells over seven days following radiation (Supp Fig-7D).

**Figure 1:**
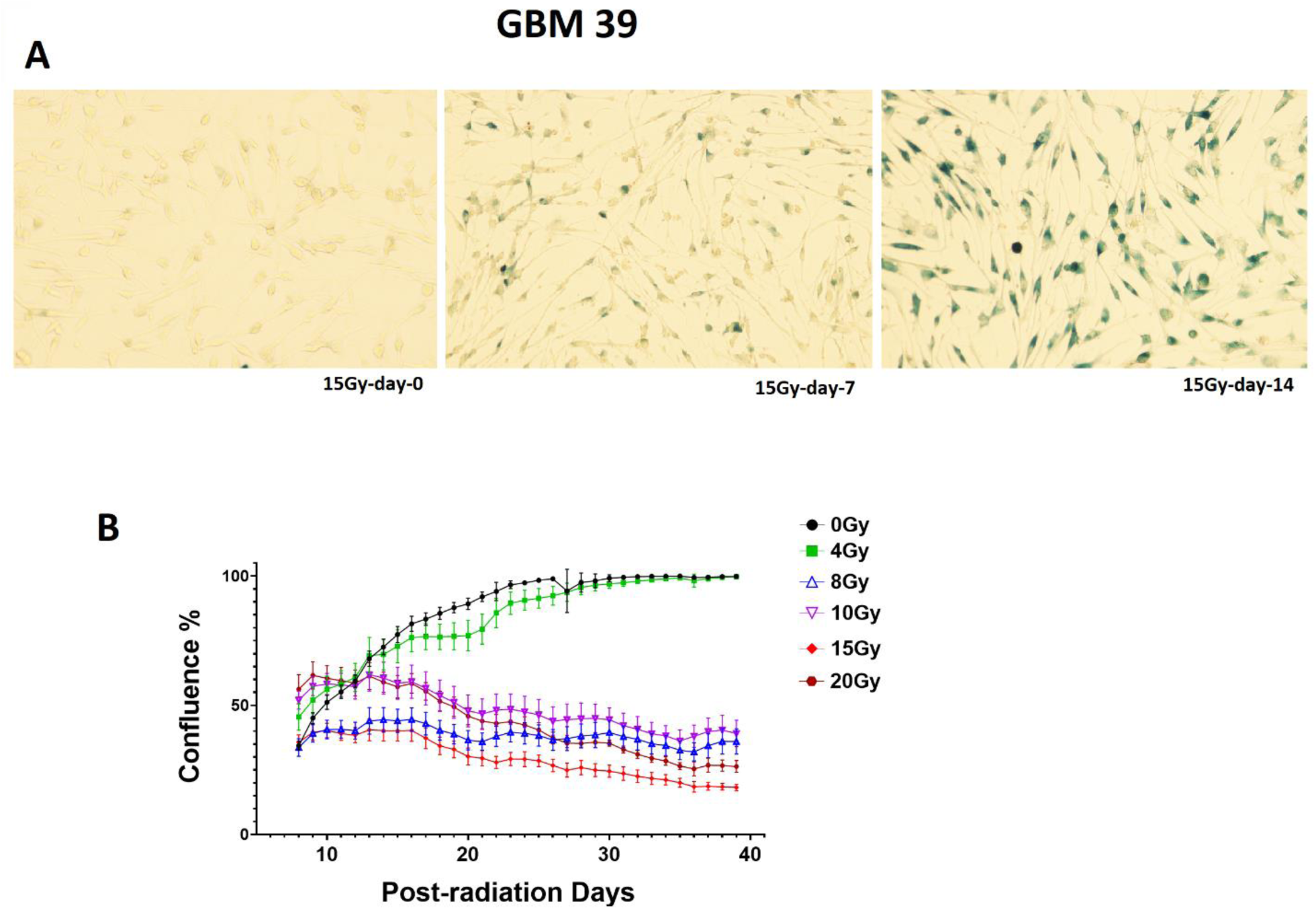
Radiation of GBM cells *in vitro* induces a senescent-like phenotype. **A:** Increased SA-Beta-Galactosidase staining in GBM 39 over 14 days following 15Gy radiation. **B:** Change in GB39 confluency *in vitro* over 40 days following radiation doses from 0-20Gy. Error bars show SEM from technical replicates.

### Radiated GBM cell lines selectively vulnerable anti-Bcl-2 family agents

A growing variety of drugs have been proposed as having senolytic activity. Consistent with certain overlapping biology between senescence and malignancy, some senolytic drugs have known anti-tumor activity.[12][14] We investigated whether prior radiation increases sensitivity to senolytic agents. The senolytic drugs tested and their associated senescent cell associated pathways (SCAPs) are detailed in Table 1. [14] Irradiated cell lines GBM39 and GBM76 both demonstrated relatively higher sensitivity to the Bcl-2 family targeting drugs A1331852 (mean +/-SD, range for each cell line) and Navitoclax (mean +/-SD, range for each cell line) than non-radiated cells (p<0.0001)(Figure 2, Supp Figure-1). Both target the Bcl-2 family of anti-apoptotic proteins. Specifically, Navitoclax targets Bcl-2 and Bcl-XL, whereas A1331852 targets Bcl-XL only.[15][16]

**Table 1.** [38]

| **Senescent Cell Anti-Apoptotic Pathways (SCAPS)** | **Inhibitors** |
| --- | --- |
| 1. PI3Kδ, AKT, ROS-protective, metabolic | Quercetin, Fisetin, Piperlongumine |
| 2. MDM2, p53, p21, serpine | Quercetin, Fisetin, Dasatinib |
| 3. Bcl-2 | Navitoclax, Venitoclax |
| 4. Bcl-XL | Navitoclax, A1331852, A1155463 |
| 5. Ephrins, dependence receptors, tyrosine kinases | Dasatinib, Piperlongumine |
| 6. HIF-1<sup>α</sup> | Quercetin, Fisetin |
| 7. HSP-90 | Onalespib |

**Figure 2:**
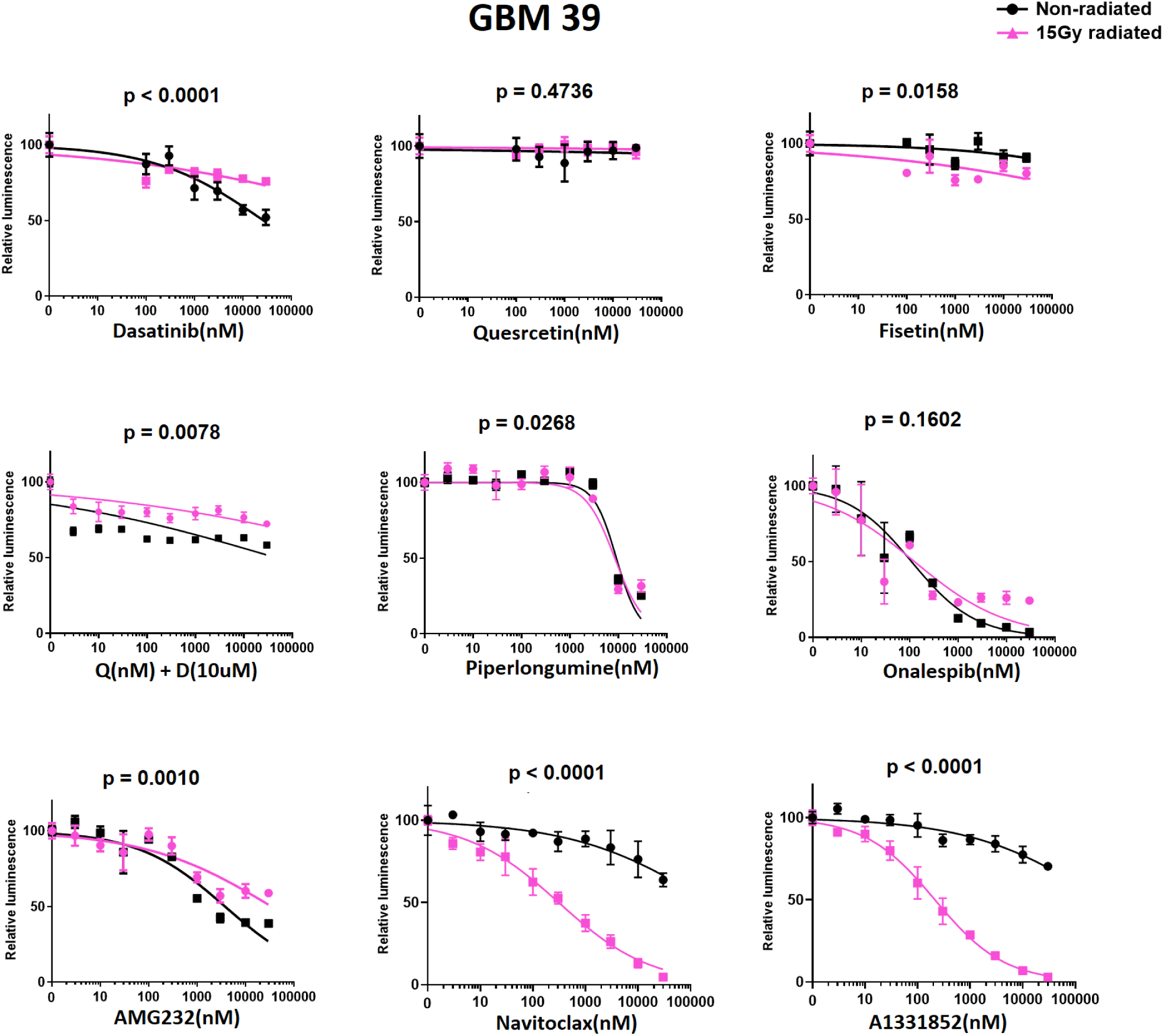
Navitoclax and A1331852 ablate radiated GBM39. Candidate senolytic drugs were evaluated using GBM39: Drug screening was performed 21 days after radiation. Cells were exposed to drugs for four days prior analysis via Cell-Titer-Glo. Purple line and black lines denote the dose-response curve for 15Gy and 0Gy radiated cells, respectively. Luminescence values are normalized to 0nM control for each radiation dose. Navitoclax and A1331852 demonstrated lower IC50 in radiated cells, p<0.0001. The data shown are means ±SEM (standard error of mean) of 3 technical replicates; similar results were obtained in GBM39 with 10 or 20Gy and GBM76 after 10, 15 or 20Gy (see Supp. fig-1)

To further discriminate between the dependence of radiated human GBM upon Bcl-XL and Bcl-2, we evaluated Navitoclax and A1331852, as well as A1155463-a Bcl-XL-selective inhibitor, and Venetoclax, a Bcl-2-selective inhibitor.[17] For reference, the previously published Ki values for each of these 4 drugs for anti-apoptotic Bcl-2 family members (Bcl-XL, BCl-2 and Bcl-W) is shown in table 4. To date, no drug is available to selectively inhibit Bcl-W. Ablation of radiated cells was seen across all lines (GBM10, 39, 76, 123), for all drugs, except for Venetoclax (Figure 3). Three of four lines showed no selective ablation with Venetoclax; lower viability was seen with high dose Venetoclax in radiated GBM 10 cells. These data suggested Bcl-XL to be the most relevant therapeutic target for ablation of previously radiated senescent-like GBM. To further evaluate the reproducibility of Bcl-XL dependence in radiated human glioblastoma, we tested A1331852 and/or Navitoclax in several additional molecularly diverse cell lines with and without prior 10, 15 or 20Gy radiation (Supp Figure-3). The molecular characteristics of the cell lines utilized are summarized in table 2. Among the 13 human GBM cell lines evaluated, meaningful analysis was possible in all with the exception of GBM43, for which insufficient cells survived radiation to permit testing. Results of the pharmacologic Bcl-XL inhibition studies performed are summarized in table 3 and Figure-7. While the IC50 of Bcl-XL inhibition with and without radiation varied markedly between lines, radiated GBM was reproducibly more susceptible to Bcl-XL inhibition than non-radiated GBM.

**Table.2.** [27]

| Line | Sex | age | Recurrence Status | MGMT methylation | Sub-type | IDH1 | CDKN2A | PTEN | EGFR | TP53 | Met | Tert Prom |
| --- | --- | --- | --- | --- | --- | --- | --- | --- | --- | --- | --- | --- |
| 6 | M | 65 | Primary | U | Clas |  | LL | L | A (v3) | M | A | M |
| 10 | M | 41 | Recurrent | U | Mes |  | LL | LM | A |  | A | M |
| 12 | M | 68 | Primary | M | Mes |  | LL |  | AM | ML |  | M |
| 39 | M | 51 | Primary | M | Mes |  | L | LM | A (v3) |  | A | M |
| 43 | M | 69 | Primary | U | Mes |  | LL |  |  | M |  | M |
| 76 | M | 38 | Recurrent | M | Clas |  | LL | LM | A (v3) |  | A | M |
| 114* | M | 68 | Primary | M | ND |  | LL | L | A |  |  | M |
| 116* | F | 56 | Primary | M | Mes |  | LL | LM | A | M | A | M |
| 120* | M | 57 | Recurrent | U | PN |  | A | LL | A | M | A | M |
| 123 | F | 62 | Recurrent | U | PN |  |  | L | A (v2) | M |  | M |
| 132* | M | 75 | Primary | U | Clas |  | L | L | A | M |  | M |
| 164 | F | 38 | Primary | M | PN | M<sup>h</sup> | LL | L |  |  | A | - |
| 196 | F | 30 | Primary | M | ND | M<sup>hh</sup> | LL | L | A | MA | A | - |

\* = dBT ("differentiated brain tumor – aka, maintained in culture).
M=Mutant
A=Amplified or net gain (>2n)
L=Loss
LL= Homozygous deletion
Subtypes: Classical (Clas), Mesenchymal (Mes), Proneural (P), Not determined (ND)
<sup>h</sup> heterozygous IDH R132H mutation
<sup>hh</sup> homozygous IDH R132H mutation
v2, v3 = EGFR variants.

**Table-3:** IC50 of A1331852(uM)

| Cell lines<br>(passage<br>number) | Control | Senescent | Relative<br>sensitivity<br>(RS) | Treatment | P value | N of<br>experi<br>ments |
| --- | --- | --- | --- | --- | --- | --- |
| 6(P4) | 14.128 | 2.26 | 6.251327434 | 10Gy | p <0.0001 | 1 |
| 6(P4) | 14.128 | 0.5316 | 26.57637 | 15Gy | p <0.0001 | 1 |
| 6(P4) | 14.128 | 0.8452 | 16.71557 | 20Gy | p <0.0001 | 1 |
| 10(P2) | 39.97 | 2.469 | 16.18874 | 15Gy | p <0.0001 | 1 |
| 12(P3) | 0.39 ± 0.07 | 0.0229 ±0.00761 | 19.7362<br>±8.282214 | 15Gy | p <0.0001 | 3 |
| 39(P2-3) | 107.8895±152.35 | 0.080547± 0.093 | 513.8149<br>±711.64 | 15Gy | p <0.0001 | 3 |
| 39(P3) | 1.30465±1.01435 | 0.071±0.010255 | 16.65831<br>±11.87971 | 20Gy | p <0.0001 | 2 |
| 39(P2-3) | 1.7164 ±2.1821 | 0.041846±2.182<br>1 | 41.01706<br>±2.182154 | TMZ(100uM) | p <0.0001 | 3 |
| 76(P2) | 23.464 ±6.762 | 0.9998 ±0.36727 | 31.77018<br>± 23.61 | 10Gy | p <0.0001 | 3 |
| 76(P2) | 23.464 ±6.7622 | 1.225 ±0.41632 | 20.64494<br>±5.632602 | 15Gy | p <0.0001 | 3 |
| 76(P2)- | 23.464<br>± 6.762161 | 0.679± 0.049209 | 34.24256<br>± 7.988135 | 20Gy | p <0.0001 | 3 |
| 76(P3)- | 1.685 | 0.05973 | 28.21028 | TMZ(100uM) | p <0.0001 | 1 |
| 114(P11)* | 6.3175 ±1.0925 | 0.33825<br>±0.07205 | 20.28±7.55 | 15Gy | p <0.0001 | 2 |
| 116(P8)* | 0.2899 | 0.06349 | 4.566073 | 20Gy | p <0.0001 | 1 |
| 120(P10)* | 5.179 | 0.4581 | 11.30539 | 15Gy | p <0.0001 | 1 |
| 123(P3) | 22.991 | 9.356 | 2.457354 | 15Gy | P =0.0532 | 1 |
| 132(P10)* | 18.166 | 4.311 | 4.213871 | 10Gy | p <0.0001 | 1 |
| 132(P10)* | 18.166 | 2.934 | 6.191547 | 20Gy | p <0.0001 | 1 |
| 164(P2) | 0.2973 | 0.07774 | 3.824286 | 15Gy |  | 1 |
| 196(P2) | 27.336 | 2.549 | 10.72420557 | 15Gy | p <0.0001 | 1 |

Relative sensitivity = fold changes of IC50 comparing to IC50 of control cells
\* = dBT (maintained in culture)

**Table 4.**
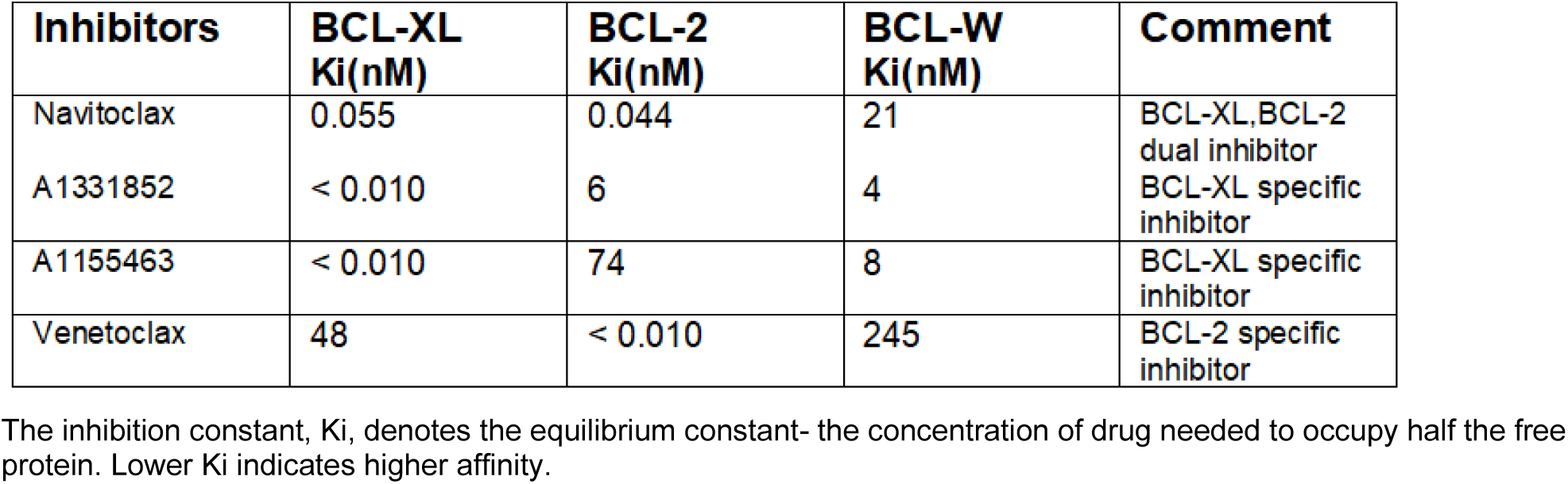
[39][40]

**Figure 3:**
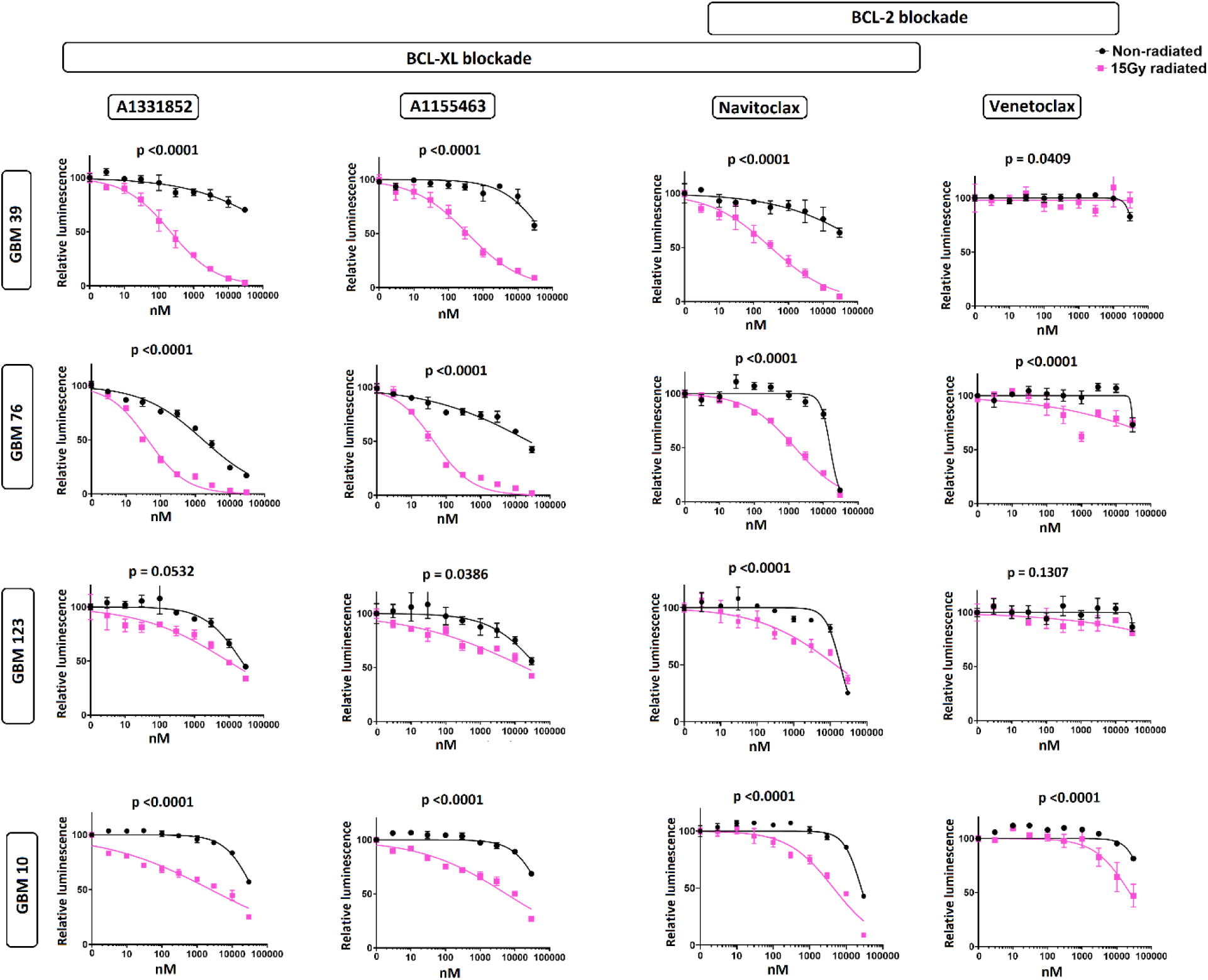
Radiated GBM cell lines are selectively vulnerable to Bcl-XL blockade. GBM39, 76, 10 and 123 were used to evaluate the senolytic activity of BCL2-family inhibitors, including the BCL-XL-specific inhibitors (A1331852, A1155463), the selective BCL-2inhibitor (Venetoclax) and dual inhibitor of both BCL-XL and BCL-2 (Navitoclax). Dose response curves shown for control non-radiated (black) and 15Gy radiated (purple) cells. Cells were exposed to drug for 4 days, starting 21 days after radiation. 15Gy-radiated cells demonstrated higher sensitivity than non-radiated cells to the BCL-X-selective inhibitors (A1331852, A1155463), and Navitoclax (Inhibits Bcl-XL and Bcl-2) but, not Venetoclax (BCL-2-selective inhibitor). For all groups, luminescence values are normalized individually to 0nM control. Data shown are means +SEM of 3 technical replicates at each concentration. Data shown are representative of multiple confirmatory experiments. Complete data for each cell line and condition are available in supplemental table-3.

**Figure 4:**
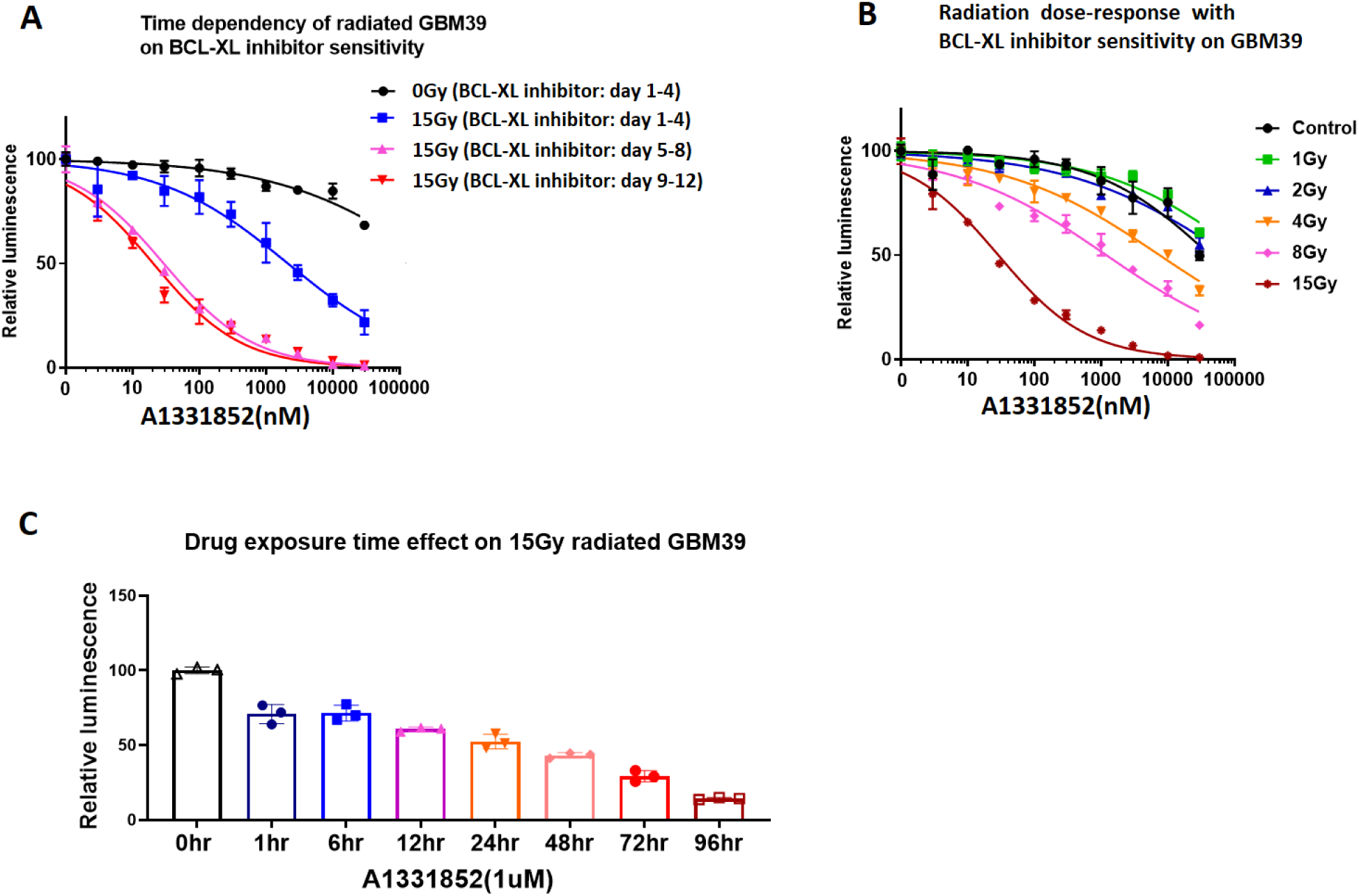
GBM vulnerability to BCL-XL inhibition depends on radiation timing, radiation dose, and duration of inhibitor exposure. **A:** Sensitivity to BCL-XL inhibition at 4 (blue), 8 (purple) and 12 (red) days after 15Gy radiation.). All cohorts were exposed to drug for the same amount of time exposure time (4 days), and were analyzed on the 5th day after plating. Very similar results were obtained with the alternate Bcl-XL inhibitor A1155463, and after induction of senescence with TMZ—see Supplementary figure 4. **B:** Impact of prior radiation dose on BCL-XL inhibitor sensitivity: A1331852 treatment was initiated 4 days following variable doses of radiation. **C:** Duration of drug exposure impacts GBM39 vulnerability to A1331852, applied 7days post radiation for 1hour to 96hours with equal total culture duration prior to analysis. Luminescence values are normalized individually by 0nM control. Graphs show means ±SEM of technical triplicates at each concentration.

**Figure 5:**
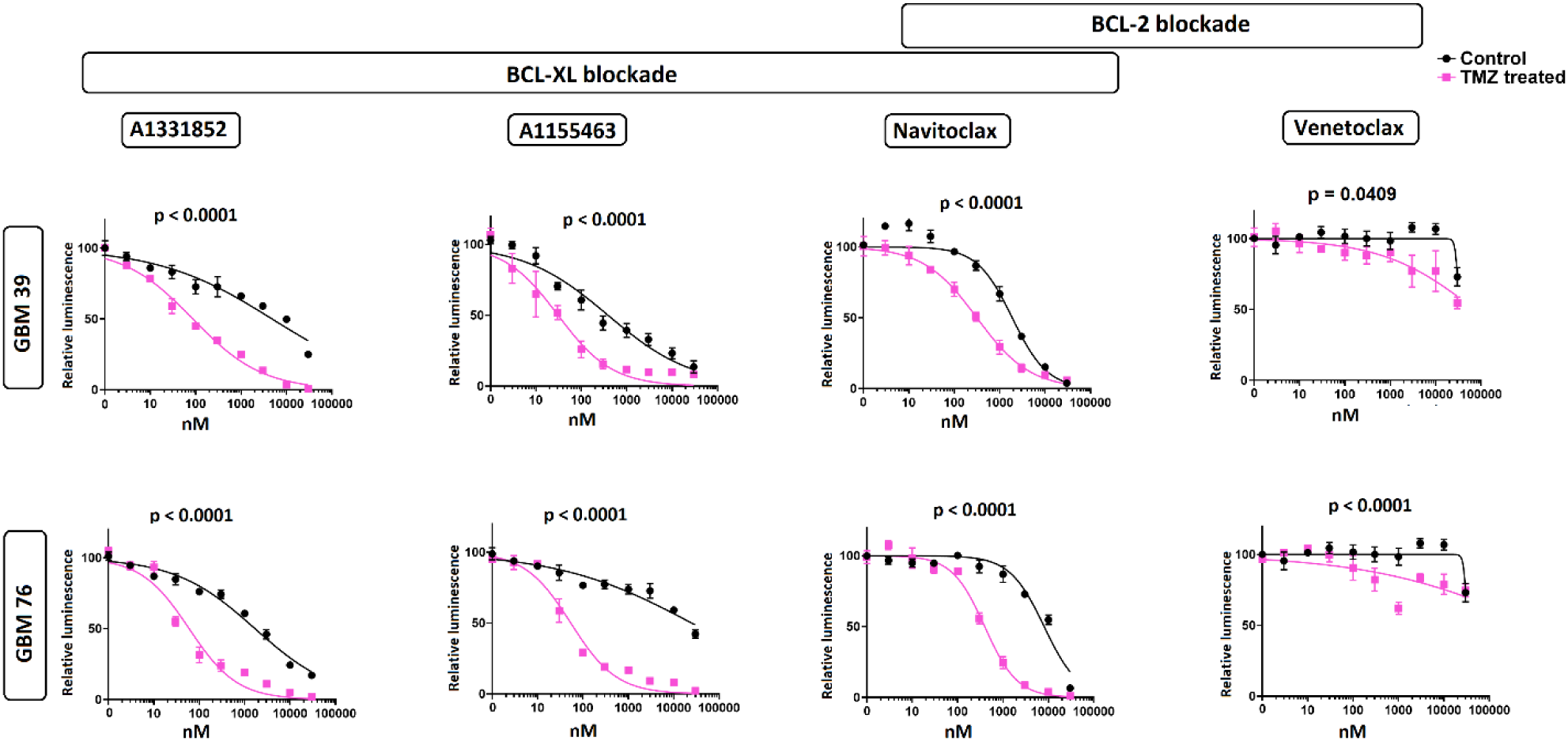
TMZ exposure induces selective vulnerability to Bcl-XL inhibitors. GBM76 and GBM39 were treated with TMZ (100uM) for 7days followed by 14 days TMZ-free media prior to treatment with Bcl-2 family inhibitors as shown. TMZ-treated cells demonstrated selective vulnerability to BCL-XL inhibitors (A1331852, A1155463, Navitoclax), but not to the BCL-2-specific inhibitor (Venetoclax). For all experiments, luminescence values are normalized individually to 0nM control. All data are means +SEM of 3 technical replicates at each concentration.

**Figure 6:**
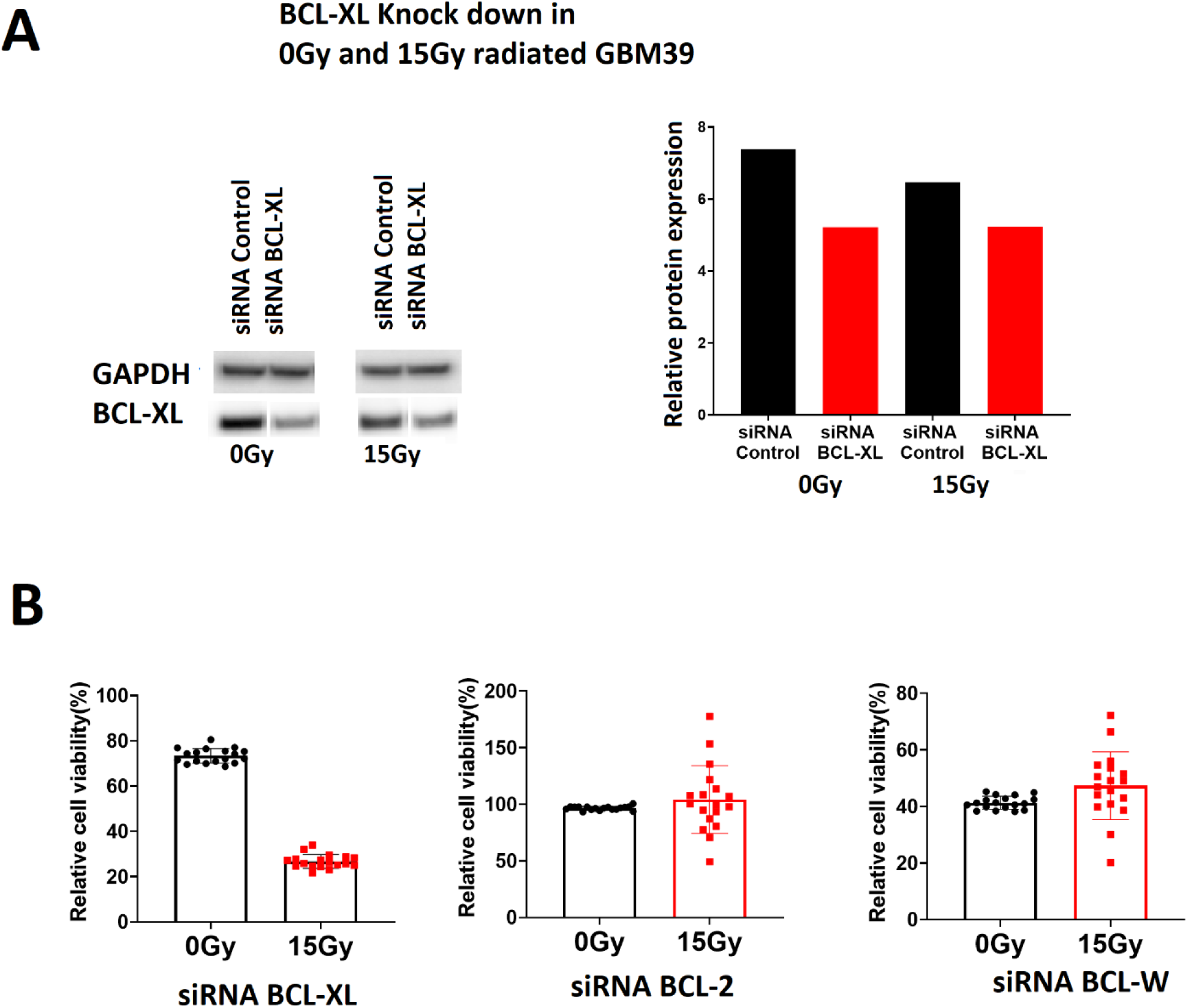
Radiated GBM is selectively vulnerable to Bcl-XL knockdown. **A:** BCL-XL, BCL-W and BCL-2 were knocked down via SiRNA in GBM 39, 7 days after 0Gy, or 15Gy radiation. Data are presented normalized to scrambled control of each group. Representative data one of three different construct of siRNA BCL-XL has shown, Western blot confirmation of knockdown following control siRNA and all constructs of siRNA BCL-XL has shown in Supp. Fig-6. **B:** Similar knockdown experiments were performed for Bcl-XL, Bcl-2 and Bcl-W. Radiated cells showed decreased survival relative to non-irradiated cells in the Bcl-XL knock down group only (p<0.0001).

**Figure-7:**
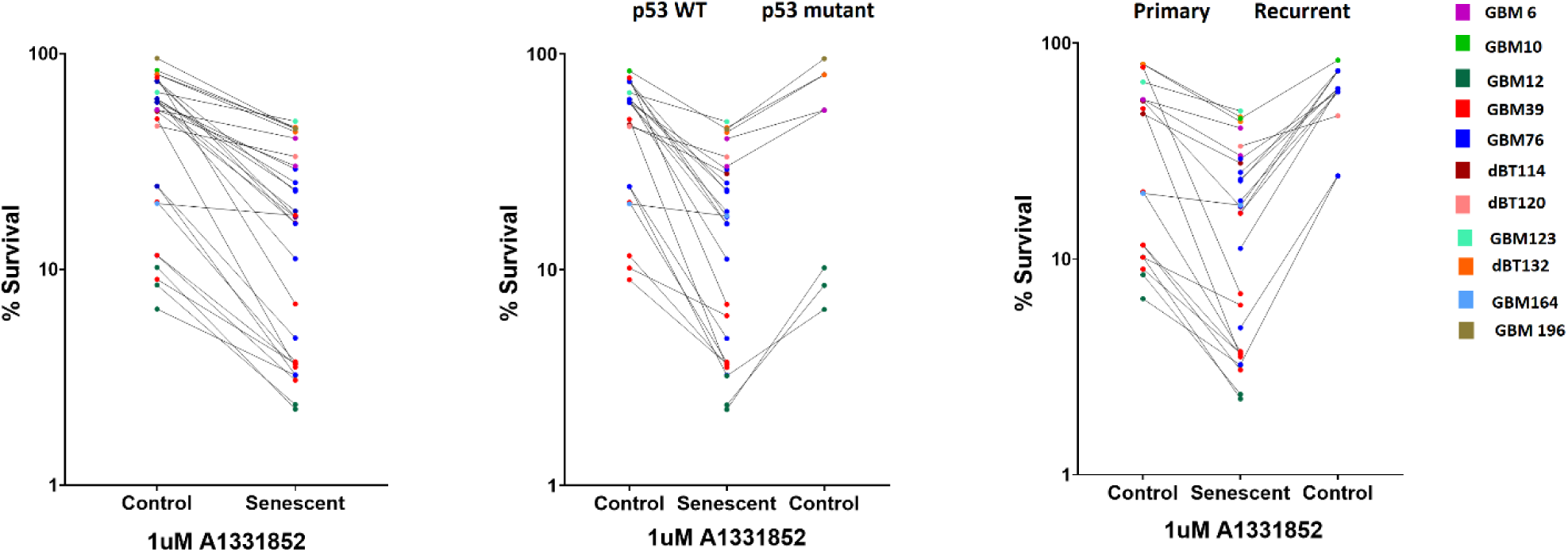
Senescent GBM is selectively vulnerable to BCL-XL inhibitors. **A:** Relative survival of senescent and proliferating all GBM cell lines tested with 1uM A1331852 treatment. **B:** Comparison of selective vulnerability of p53 wild type and mutant senescence Glioma to BCL-XL inhibition. C: Demonstrating senolytic effect of A1331852 across the primary and recurrent Glioma with or without Chemoradiation.

### Time-point of BCL-XL inhibitor sensitivity

The previous experiments maintained radiated cells for 2-4 weeks after radiation prior to senolytic drug testing. While the senescence process often takes weeks, the apoptotic pathways regulated by bcl-2 family members are dynamically regulated within days following DNA damage. As such, we used GBM39 to ask if a minimal period of time after radiation must elapse following radiation prior to onset of Bcl-XL sensitivity. Using a timed assay following radiation, GBM39 was treated with Bcl-XL inhibitors A1331852 or A1155463 for 4 days, starting 1, 5, or 9 days after radiation. Analysis was performed at the end of the 4 days drug exposure. While efficacy was seen upon treatment starting one day after radiation, more complete cell ablation was observed with a leftward shift of the dose response curve, four days elapsed after radiation prior to starting treatment 5 or 9 days following radiation (Fig 4A and Sup Fig 4B).

### Dependency on drug exposure time and radiation dose

We next asked if a minimal radiation dose was required to induce susceptibility cell death with Bcl-XL inhibition. Radiation doses of 4Gy or higher in GBM39 promoted sensitivity to A1331852 with increasing efficacy up to 15Gy (Fig 4B). For most cell lines, radiation doses of 10-20Gy yielded similar results (Sup fig 3, and Table 3). In anticipation of future considerations related therapeutic dosing, we next asked what duration of continuous BCL-XL inhibitor exposure may be required to observe senolytic effect. To accomplish this, GBM39 was exposed to drug for varying durations from 0-96 hours, with drug then washed off and cells maintained in normal growth media thereafter until time of analysis. Although some most impact was observed even with 1hour of treatment, maximal impact was seen in cells that received sustained exposure to Bcl-XL inhibition for 96hours.

### Elimination of TMZ treated senescent glioma

Radiation and TMZ are both routinely administered to patients with GBM. Both may induce senescence and modulate apoptotic machinery. To determine whether TMZ (100uM) exposure induces selective susceptibility to BCL-XL inhibition, we pre-treated the GBM cell lines with TMZ for 20 days and then analyzed various anti-Bcl-family agents. Using GBM76, and GBM39, we found that prior 20 days TMZ exposure induced sensitivity to the BCL-XL inhibition, but not BCL-2 inhibition as previously observed following radiation (Figure 4 and Supp Figure 4).

Based on the previously published selectivity of A1331852, A1155463 to Bcl-XL and Venetoclax to Bcl-2 (table 4), we predicted that knockdown of Bcl-XL but not Bcl2 would impede survival in radiated cells. We utilized siRNA constructs and scrambled controls and evaluated their impact on survival in radiated and non-radiate GBM. Compared to scrambled controls, Bcl-2 and Bcl-W knockdown elicited no impact on cell survival whereas Bcl-XL knockdown significantly decreased survival of radiated cells.

Finally, we investigated whether prior cytotoxic therapy would be sufficient to permanently induce sensitivity to Bcl-XL blockade, or if cells must remain in a senescent-like non-proliferative state to maintain sensitivity. In one culture maintained over 6 weeks following 8Gy radiation, proliferative activity resumed, with a doubling time ultimately unchanged from the parent culture. Cells in the culture that “escaped senescence” to resume proliferative behavior lost sensitivity to Bcl-XL inhibition (Supp Figure 5). This result is consistent with the observation that gliomas derived from both primary and recurrent lesions (Supp Figure 3) proved relatively insensitive to Bcl-XL inhibition until placed into a state of proliferative arrest with either TMZ or radiation.

## Discussion

We here demonstrate that human GBM cells surviving in a senescent-like state after chemotherapy or TMZ are relatively more susceptible to BCL-XL-blockade than naïve proliferative GBM. Conventional cancer therapies such as radiation and alkylating chemotherapies act through induction of oxidative stress and DNA damage. [18][19] These can exert therapeutic impact through multiple mechanisms: (i) lethal cell damage (ii) inducing a senescent-like state of proliferative arrest that impedes further growth,[20] (iii) promoting a state of tumor quiescence to attenuate proliferation, and/or (iv)altering the tumor microenvironment or tumor-immune response to indirectly achieve some combination of the first three goals.[21]

That gliomas invariably recur highlights the inadequate levels of lethal cell damage achieved with standard chemoradiation. Gliomas are well described to harbor a population of relatively quiescent and resilient glioma stem cells.[7][22][23] Some have interpreted glioma recurrence as evidence that quiescent “glioma stem cells” persist in a quiescent state after chemoradiation prior to ultimately re-initiating tumor recurrence.[24] Other evidence in glioma and other tumors supports induction of a senescent-like state, after cytotoxic therapy. If indeed quiescent glioma stem cells and senescent cells *both* exist in glioma after standard therapy, do they both actively coexist in parallel at all points following therapy, or could initially senescent cells escape senescence to revitalize the tumor as quiescent stem cells? Recent work showed that that cancer cells with an initially senescent phenotype following chemotherapy escaped senescence upon p53 inhibition giving rise to tumor stem cells that were more aggressive and resilient than those present in the original tumor.[25][26] If true for glioma, senolytic therapies could help eliminate senescent glioma cells before they have the chance to re-emerge as quiescent glioma stem cells. Indeed, we found that in one culture of GBM39 that eventually re-entered cell cycle 6 weeks after 8Gy, proved highly resistant even to unable to promote apoptosis, even at doses that had been effective prior to the original radiation.

Conversely, four of the GBM lines used (10, 76, 120, 123) were derived from patient with recurrent tumors who had previously undergone chemotherapy and radiation.[27] Although GBM123 was among the weaker responders, all others showed similarly augmented response to BCL-XL inhibition after re-induction of senescence with radiation or TMZ. Also of note, certain lines tested (including GBM 6 and 10) have been previously reported as radio-resistant with nominal prolongation of survival in xenografts upon radiation. Nevertheless, these were amenable to sufficient proliferative arrest in vitro, to permit augmented ablation with BCL-XL inhibition.

p53 is a key regulator of cell cycle arrest in the context of radiation-induced DNA damage and senescence, and is the most commonly mutated gene across human cancers.[28][29] Although several lines with some of the most robust relative sensitivity to BCL-XL (in radiated vs non-radiated cells) were WT for p53 (e.g., GBM 10, 39, 76, 114), highly significant induction was also observed in p53-mutant cell lines (GBM 6, 12, 132, 196). Similarly, no clear p53 pattern followed for cells with modest radiation-induced induction BCL-XL inhibitor sensitivity: P53-mutant GBM123 showed only 2.5x induction of A1331852 sensitivity (results were more robust for A1155 and Navitoclax, Figure 3), yet p53-WT GBM 164 showed minimally significant difference between radiated and non-radiated cells for both A133A852 and Navitoclax. That 11 of 12 tested lines showed significant induction of sensitivity to one or more BCL-XL inhibitors with radiation supports the broadly reproducible phenomenon across molecularly diverse GBM

It is likely that additional stimuli besides radiation and alkylating chemotherapy may augment sensitivity to BCl-XL inhibition. Massler, *et* al., previously reported induction of synthetic lethality with BCL-XL inhibition in IDH-mutant gliomas, or upon induction of the on cometabolite D2-HG produced by IDH-mutant gliomas. [30] Their study specifically utilized GBM164, which they observed to have a robust sensitivity to BCL-XL. Indeed, among the 12 lines tested, GBM164 was one of the two most sensitive prior to radiation. However, the most sensitive line was GBM 12, which is IDH wild-type. We additionally evaluated another line which unlike GBM196, which is homozygous for IDH1-R132H, in contrast to GBM164 which is heterozygous for IDH1-R132H. Accordingly, GBM196 generates higher levels of D2-HG (not shown), yet demonstrated ∼100x lower baseline sensitivity to A1331852 than GBM164. While we did not independently evaluate the impact of D2-HG on BCL-XL inhibition in this study, data obtained in this study would have not provided a rationale to do so. Rather, our data indicate that baseline sensitivity to BCL-XL inhibition may vary widely among both IDH-mutant and IDH-WT cell lines. However, in 90% of cases, radiation exposure was observed to enhance radiation sensitivity. It should also be noted that baseline sensitivity to BCL-XL was not predictive of the degree of final BCL-XL sensitivity achievable after exposure to radiation or TMZ. Indeed, certain lines with negligible baseline sensitivity (e.g. GBM39) were induced by radiation to be among the most sensitive after radiation.

As such, while some molecular therapies can be selectively targeted to specific tumor subtypes, we don’t our current data with only 12 cell lines divergent across phenotypes for EGFR, PTEN, CDKN2A, P53, as well as gender, age, MGMT methylation, molecular subtype and recurrence status revealed no specific molecular phenotype that would obviously portend poor response to Bcl-XL inhibition. Rather, we speculate that the therapy-induced sensitivity to Bcl-XL dependency may be mechanistically linked to intracellular processes subserving and maintaining a molecular mediator of proliferative arrest, and could thus be synergistic with therapies that activate appropriate mitotic checkpoint machinery. [31] Among master regulators of cellular senescence, a mechanistic role of p16 is largely excluded by virtue of most tested lines being homozygous null for CDKN2A, which encodes p16, an inhibitor of CDK4/6.[32][33][34] This could by implication reduce the probability of synergy between Bcl-XL inhibitors with CDK4/6 inhibitors, though also induce senescence, at least in regards to Bcl-XL inhibition. Conversely, p21 is almost ubiquitously conserved in glioma, is frequently overexpressed in glioma and is upregulated by radiation-induced DNA damage, providing an attractive avenue for future mechanistic investigations. While P21 is upregulated by p53 which is mutant in many GBMs, p21 is also regulated by other pathways and the interactions between mutant p53 and p21 can be paradoxical in cancer.

Recent work demonstrated that systemic Navitoclax was effective to prolong survival of GBM164 in a xenograft model.[30] If BBB penetration were equivalent across cell lines, this would suggest that GBM 114, 116, 120 and 76, 39, and 12 should exhibit comparable or better sensitivity after chemotherapy and/or radiation, whereas even radiated GBM 132, 10 and 123 and 196 each maintained an in vitro IC50 >10x that of non-radiated GBM164 and may thus be more resistant. Efficacy of Navitoclax as a senolytic was recently demonstrated in a tauopathy model of Alzheimer’s disease suggesting that BBB disruption may not be required for efficacy.[35] Moreover, to the extent that radiation-induced senescence may contribute to radiation-induced cognitive impairment, concurrent ablation of senescent host cells could provide a cognitively beneficial adjunct to senescent tumor ablation.[36]

While senescence serves to attenuate cell-intrinsic proliferation, SASP factors produced by senescent cells have been shown to promote tumor infiltration and recruitment of pro-tumorigenic tumor-associated macrophages. We and others have observed increased aggressiveness of human glioma implanted into the previously radiated brain. Whether this phenomenon could be attenuated with senolytics is under investigation. However, the phenomenon of “senescent spreading” has been proposed as a means via which senescent cells could actually attenuate proliferation of adjacent tumor cells. The diversity of responses to BCL-XL inhibition in absence of clear phenotypic correlates suggest empiric studies will be required to discriminate among these possibilities.

The present study provides our first attempt to harness the senescent-like biology of treated human GBM to facilitate the clearance of dormant cells. An obvious limitation of this work is that in vitro cultures offer notoriously poor surrogates of the in vivo human glioma microenvironment. A dedicated effort to rigorously characterize the senescent tumor is of high importance. Efforts to do so for human glioma may be hampered by the rarity of surgery for dormant disease, as well as the likely paucity of the offending latent cells, as well as inherent inability to expand senescent cells. As such best available models will be needed to determine if the senescent-like state of sustained proliferative arrest seen vitro, might be antagonized by complex in vivo biology. For example, tumor-associated hypoxia in vivo with its relative nutrient deprivation may attenuate mTOR signaling which is important to achieve a stably senescent cell phenotype—Bcl-XL-related implications of which are unclear. Moreover, the complex tumor microenvironment will predictably endow malignant cells with diverse sources of trophic support that are lacking in vitro potentially requiring adjuvant strategies not anticipated from in vitro analyses.[37]

In conclusion, we demonstrate that radiation and TMZ endow glioma with increased dependence upon Bcl-XL, blockade of which may facilitate ablation of latent glioma cells that survived prior cytotoxic therapy. Since radiation and TMZ are standard of care for GBM, further work is needed to determine if Bcl-XL inhibition could be leveraged to help forestall glioma recurrence.

---

**Supplementary Figure 1:**
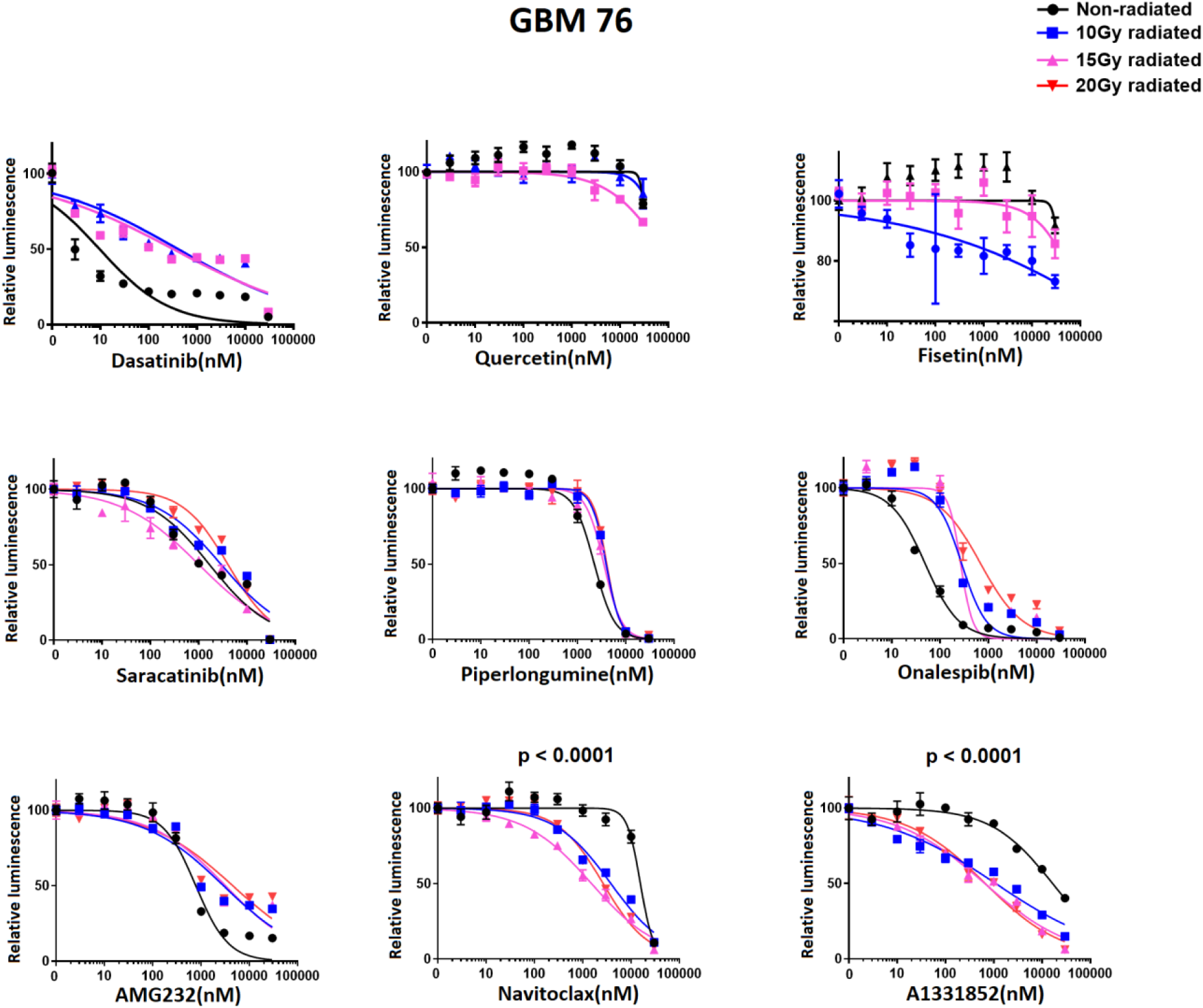
Navitoclax and A1331852 ablate radiated GBM 76 (continuation of main figure 2, which shows data for GBM39). Senolytic drug screening in GBM76 cell line with senolytic drugs targeting different anti-apoptotic pathways, Piperlongumine, MDM2 inhibitor (AMG-232), Onalespib, Dasatinib, Quercetin Fisetin, Navitoclax, A1331852 and Saracatinib. Cells were exposed to ten different concentrations of drug for 4days. Black line denotes non radiated control cells, blue line denotes 10Gy, purple line denotes 15Gy and red line denotes 20Gy radiated cells. Luminescence values are normalized individually by 0nM control. All the data are means ±SEM of triplicates at each concentration. Navitoclax and A1331852 have shown lower IC50 radiated cells comparing to non-radiated control.

**Supplementary Figure-2:**
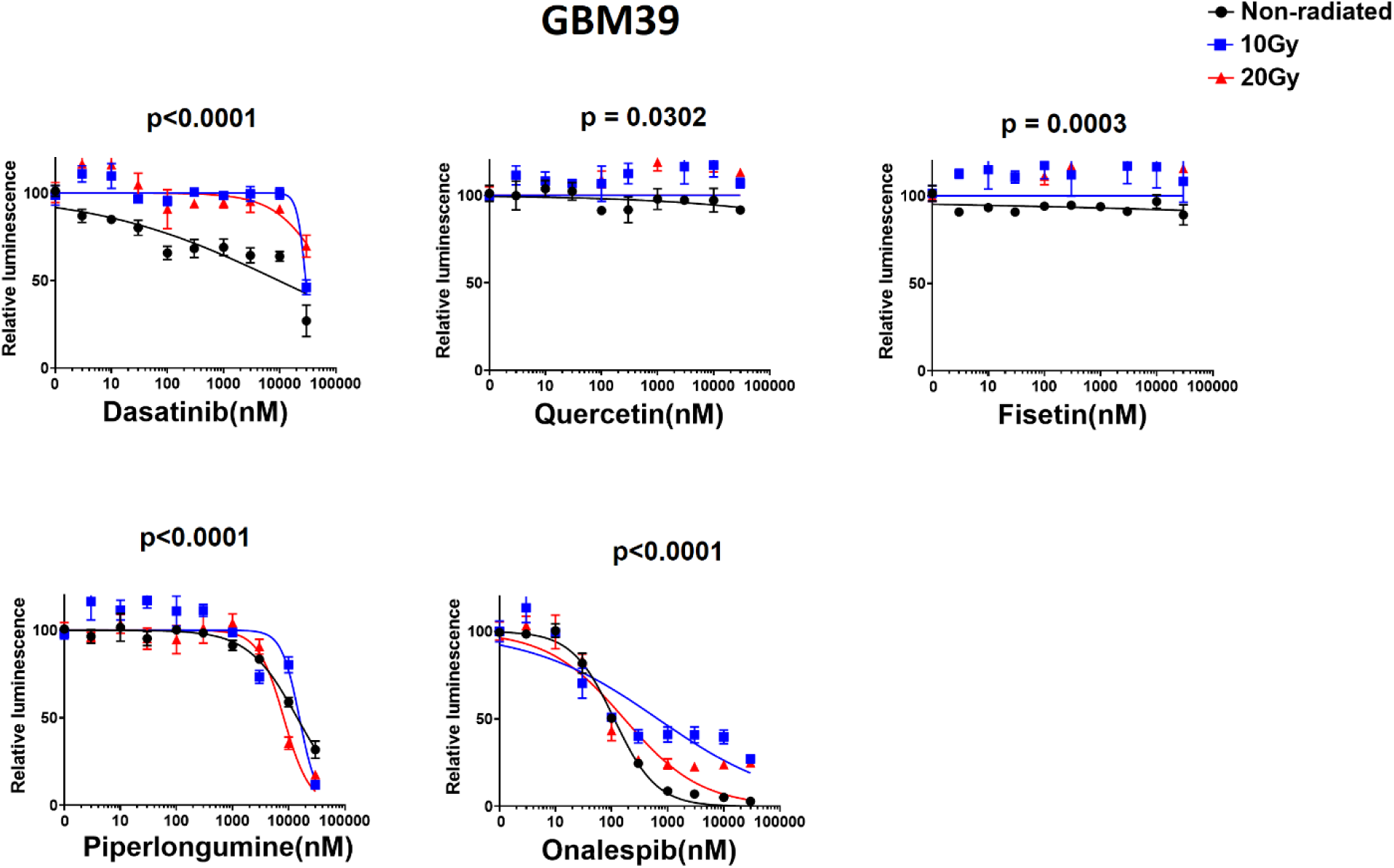
Senolytic Candidates evaluated using GBM39. (Continued from Main Figure 2), which shows data for 15Gy. Candidate senolytic drugs were evaluated using GBM39 with and without prior exposure to 10 or 20Gy. Black line denotes non radiated control cells, blue line denotes 10Gy, and red line denotes 20Gy radiated cells. Luminescence values are normalized individually for each radiation dose to the 0nM control. All the data are means ±SEM of triplicates at each concentration. Navitoclax and A1331852 have shown lower IC50 radiated cells comparing to non-radiated control.

**Supplementary figure-3:**
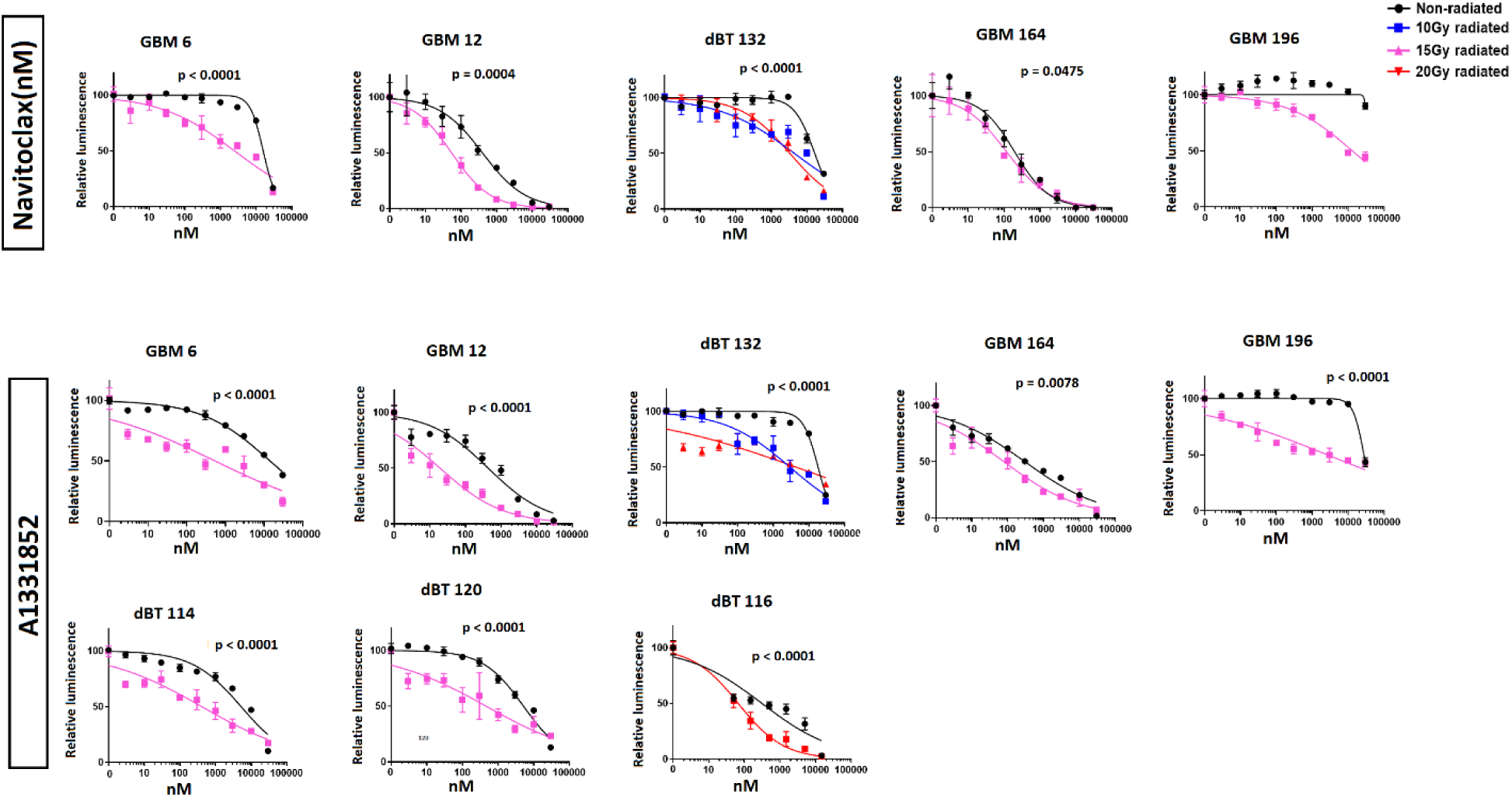
Multiple radiated GBM cell lines are selectively sensitive to Bcl-XL inhibition. All cell lines tested demonstrated increased sensitivity to BCL-Xl inhibition after prior radiation. For all experiments, luminescence values are normalized to the 0nM control for that cell line and radiation dose. All data are means +SD of 3 technical replicates at each concentration.

**Supplementary figure-4:**
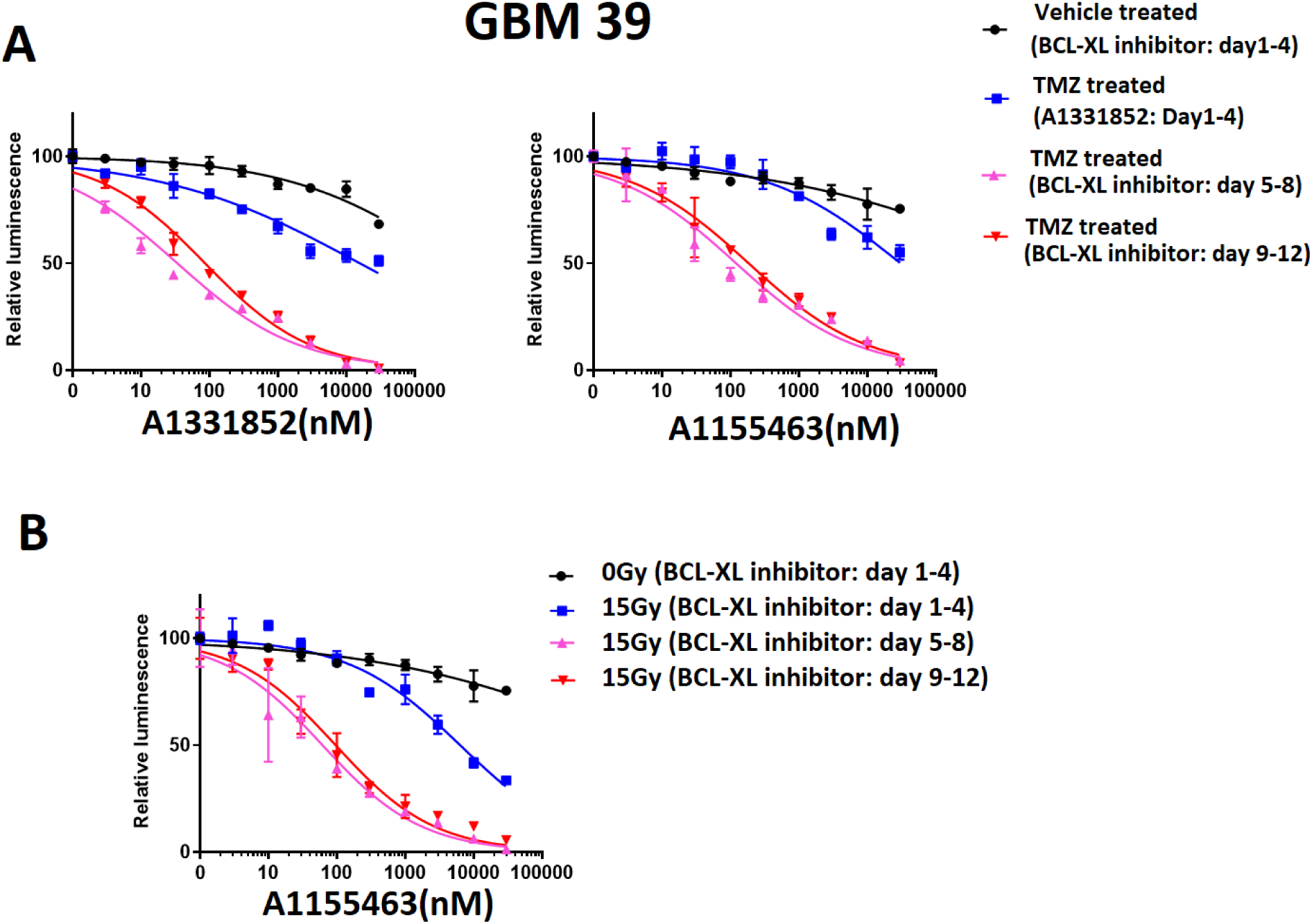
Navitoclax and A1331852 in TMZ-treated GBM39 (continued from Main Figure 4a): Evaluation of time-dependency of glioblastoma for BCL-XL inhibitor-mediated ablation. **A:** Sensitivity to BCL-XL inhibitors (A1331852 and A1155463) at different time-points following TMZ treatment; blue line denotes 4days post TMZ, purple line denotes 8 days post TMZ and red line denotes 12 days post TMZ treatment. **B:** Sensitivity to BCL-XL inhibitors, A1155463 at different time-points following 15Gy radiation. Blue line denotes 4days post radiation, purple line denotes 8 days post radiation and red line denotes 12 days since radiation with 4 days drug exposure to all cohorts. Luminescence values are normalized individually by 0nM control. All the data are means ±SEM of triplicates at each concentration.

**Supplementary figure-5:**
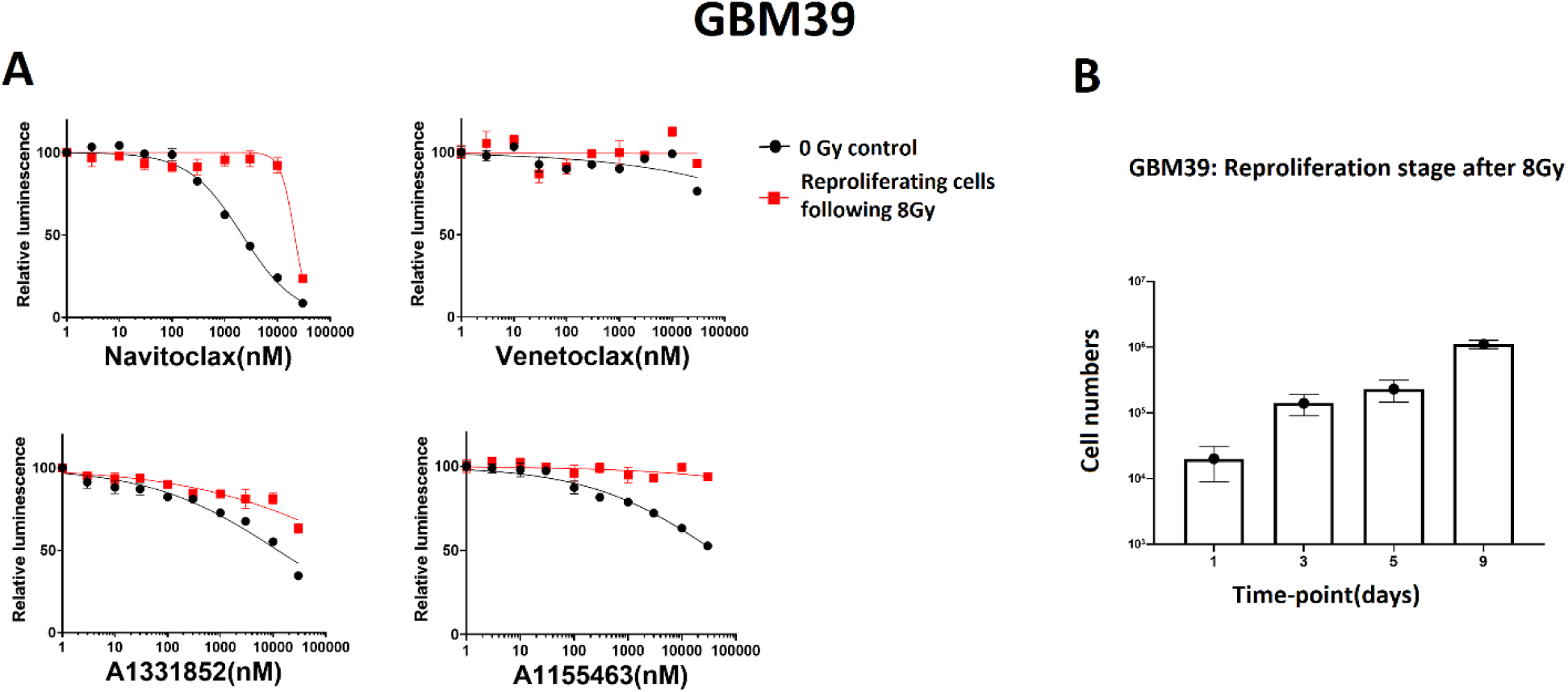
Previously radiated GBM39 fails to demonstrate sensitivity to Bcl-XL inhibition after re-entering cell cycle. **A:** BCL-2 family inhibitor drug screening in 8Gy GBM39, those restarted cell proliferation after 6weeks following radiation. Re-proliferation was detected by regular microscopic evaluation, which evident by turning into confluent from 60% post radiation confluency over 6weeks. **B:** Increasing number of cells over 9days after regaining proliferation. Cells were re-plated with equal density into 12 wells plate (day 0), and cell counts performed at day1,3,5,9. Data represented the mean± SD of three technical replicates. Drug screening (A) was performed using the cells harvested at day 5 and 9. Both demonstrated similar results, here data are shown from day 9.

**Supplementary figure-6:**
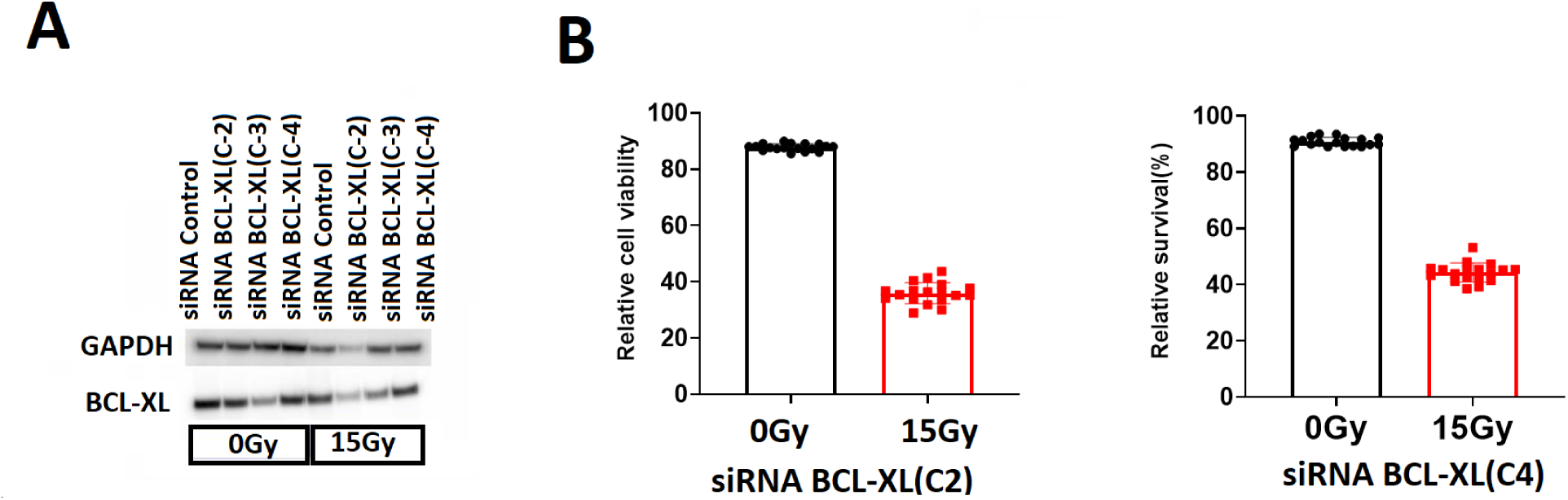
Radiated GBM is selectively vulnerable to Bcl-XL knockdown. (Continued from main figure 6): BCL-XL protein expression in 0Gy and 15Gy radiated GBM39. **A:** Western blot confirmation of knockdown following control siRNA and three different constructs of siRNA BCL-XL. **B:** Relative cell survival following BCL-XL knockdown has shown 15Gy radiated cells are more dependent than 0Gy cells upon BCL-XL for survival.

**Supplementary figure-7:**
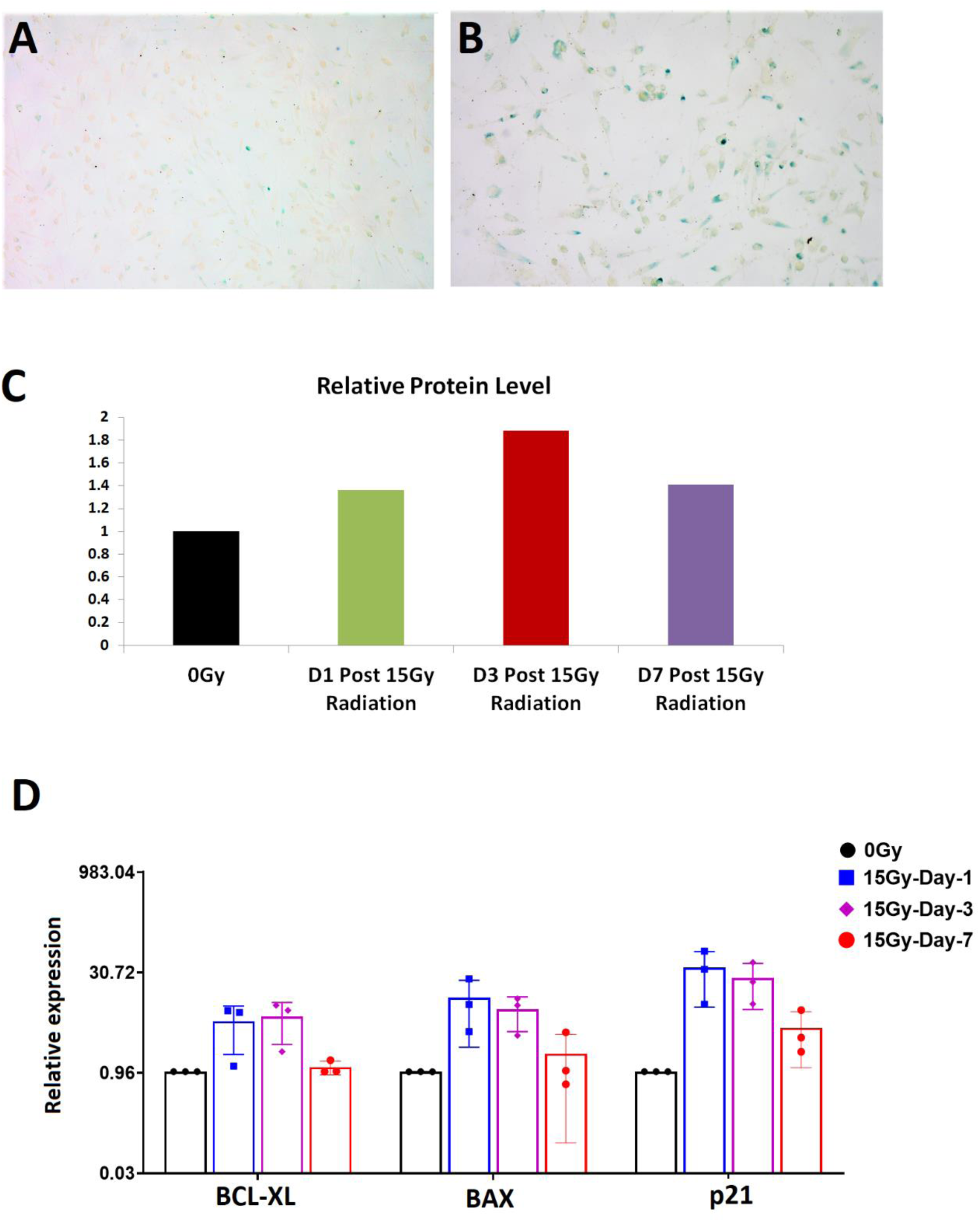
Confirmation of therapy associated senescence in GBm39. **A, B:** SA-beta gal staining of control, and TMZ (100uM) treated cells demonstrating presence of senescent glioma in treatment group. **C:** Western blot analysis showing BCL-XL expression in 0Gy and 15Gy radiated GBM39 over 7days post-radiation. **D:** qRT-PCR of GBM39 following 15Gy radiation in vitro demonstrates increasing expression of senescence-associated transcripts, p21 and anti-apoptotic BH3 family member Bcl-XL over 7 days following radiation.

